# Production of anti-SARS-CoV-2 hyperimmune globulin from convalescent plasma

**DOI:** 10.1101/2020.11.18.388991

**Authors:** Peter Vandeberg, Maria Cruz, José María Diez, W. Keither Merritt, Benjamin Santos, Susan Trukawinski, Andrea Wellhouse, Marta José, Todd Willis

## Abstract

**BACKGROUND:** In late 2019, the SARS-CoV-2 virus emerged in China and quickly spread into a world-wide pandemic. Prior to the development of specific drug therapies or a vaccine, more immediately available treatments were sought including convalescent plasma. A potential improvement from convalescent plasma could be the preparation of anti-SARS-CoV-2 hyperimmune globulin (hIVIG).

**STUDY DESIGN AND METHODS:** Convalescent plasma was collected from an existing network of plasma donation centers. A caprylate/chromatography purification process was used to manufacture hIVIG. Initial batches of hIVIG were manufactured in a versatile, small-scale facility designed and built to rapidly address emerging infectious diseases.

**RESULTS:** Processing convalescent plasma into hIVIG resulted in a highly purified IgG product with more concentrated neutralizing antibody activity. hIVIG will allow for the administration of greater antibody activity per unit of volume with decreased potential for several adverse events associated with plasma administration. IgG concentration and IgG antibody specific to SARS-CoV-2 were increased over 10-fold from convalescent plasma to the final product. Normalized ELISA activity (per mg/mL IgG) was maintained throughout the process. Protein content in these final product batches was 100% IgG, consisting of 98% monomer and dimer forms. Potentially hazardous proteins (IgM, IgA, and anti-A, anti-B and anti-D antibodies) were reduced to minimal levels.

**CONCLUSIONS:** Multiple batches of anti-SARS-CoV-2 hyperimmune globulin (hIVIG) that met regulatory requirements were manufactured from human convalescent plasma. The first clinical study in which the hIVIG will be evaluated will be Inpatient Treatment with Anti-Coronavirus Immunoglobulin (ITAC) [NCT04546581].

## Introduction

The SARS-CoV-2 virus emerged in late 2019 from the Wuhan region of China^1^, and by early 2020 the resulting infection, termed COVID-19, had reached pandemic status according to the World Health Organization.^2^ The novelty of this pathogen prompted an urgent search for therapeutic agents with efficacy against SARS-CoV-2. Convalescent plasma was quickly identified and investigated as a potential treatment for COVID-19.^3^

Convalescent plasma has a long history of treatment of infectious diseases extending from the Spanish flu pandemic^4^ to more recent outbreaks of severe acute respiratory syndrome (SARS)^5^, and Middle East respiratory syndrome (MERS).^6^ There are, however some disadvantages of convalescent plasma, including that the nature, titer and neutralizing power of the antibodies therein can vary greatly from one donor to another.^7^

Many of these disadvantages can be overcome through the purification and concentration of the specific antibodies into drug preparations. Human immunoglobulin products made from pooled plasma of convalescent or vaccinated donors have been used for many years in the treatment of various infections. Here we report on collection of convalescent plasma and manufacture anti-SARS-CoV-2 hyperimmune globulin (hIVIG) for use in clinical studies.

## Materials and Methods

### Collection of Convalescent Plasma

Plasma was collected from COVID-19 convalescent donors using a network of plasma donation centers geographically dispersed throughout the United States (Grifols network: Biomat USA Inc., Interstate Blood Bank Inc., and Talecris Plasma Resources, Inc.). All donations were collected by plasmapheresis and met the requirements for source plasma which includes screening for a number of common viruses.^8,9^ All donors were required to have laboratory evidence of COVID-19 infection, either through nucleic acid amplification testing (NAT), positive antigen test, or by SARS-CoV-2 antibody test prior to enrollment. Symptomatic donors must have complete resolution of symptoms at least 14 days before donation if they were negative by follow-up NAT, or 28 days if they had no follow-up test. Similarly, asymptomatic donors (positive by NAT or antigen tests) must wait 14 days after the initial test if they had a follow-up negative NAT but must wait 28 days after the initial test if they had no follow-up test. Asymptomatic donors tested only by an anti-SARS-CoV-2 antibody test were required to wait seven days prior to donation, but could donate immediately if they had a negative NAT. Donors also had to be negative for human leukocyte antigen (HLA) antibodies.^10^ All donations were tested by NAT (Procleix SARS-CoV-2 assay, Grifols Diagnostic Systems, Emeryville, CA) to confirm absence of SARS-CoV-2 infection and by serology to confirm presence of anti-SARS-COV-2 IgG (Architect SARS-CoV-2 IgG assay, Abbott Ireland, Diagnostics Division, Finisklin Business Park, Sligo, Ireland or Recombivirus Human Anti-SARS-CoV-2 Virus Spike 1 IgG ELISA Kit, Alpha Diagnostic International, San Antonio, TX).

### Manufacture of Convalescent SARS-CoV-2 Immune Globulin, Human

Prior to pooling, plasma pools were modeled to maintain distribution consistent with overall donor ABO blood type distribution. This maintained consistent batch to batch levels of anti-A and anti-B. Type O and Type B donors were limited to no more than two units from any single donor to decrease the likelihood of having high anti-A titers in the final product.

Pooled convalescent plasma was subjected to alcohol fractionation and caprylate/chromatography purification, resulting in highly purified IgG solutions formulated at 10% protein with glycine at a low pH. These production processes, including formulation, are identical to those for Gamunex immune globulin.^11,12^ These processes include multiple steps validated for the removal and/or inactivation of viruses.^13^

### Characterization of hIVIG Product

#### Routine batch testing

Routine batch quality control testing of hIVIG includes several measurements to characterize the product and ascertain that it is suitable for use. This characterization includes analyses for glycine, pH, protein concentration, osmolality, composition by electrophoresis, molecular weight profiling by size exclusion chromatography and anticomplement activity. Analyses are also performed for sodium caprylate, residual IgA and IgM, prekallikrein activator (PKA), factor XIa, anti-A, anti-B, and anti-D. These tests have recently been described for other immune globulin products manufactured with the caprylate/chromatography process.^12^ In addition, compendial tests for sterility and pyrogenic substances are performed on all batches.^14,15^

#### Anti-SARS-CoV-2 ELISA

Anti-SARS-CoV-2 IgG titers were determined using Human Anti-SARS-CoV-2 Virus Spike 1 (S1) IgG assay (Alpha Diagnostic). hIVIG batches were tested using multiple serial dilutions and a curve constructed by plotting the log of the optical density as a function of the log of the dilution. The titer was defined as the dilution at which this curve is equal to the low kit standard. The titer is also expressed as a ratio to an in-house control, which consists of a commercially available chimeric monoclonal SARS-CoV-2 S1 antibody (Sino Biologicals, Beijing, China) spiked into plasma from non-COVID-19 donors at levels intended to give titers similar to those found in plasma of COVID-19 donors.

#### Anti-SARS-CoV-2 Neutralizing antibody assay

Multiple batches of hIVIG were tested for anti-COVID-19 antibodies using an immunofluorescence-based neutralization assay performed at the National Institutes of Health Integrated Research Facility, Frederick, MD. This assay quantifies the anti-SARS-CoV-2 neutralization titer using a dilution series of test material tested for inhibition of infection of cultured Vero (CCL-81) cells by SARS-CoV-2 (Washington isolate, CDC) at a multiplicity of infection of 0.5. Potency was assessed using a cell-based immunosorbent assay to quantify infection by detecting the SARS-CoV-2 nucleoprotein using a specific antibody raised against the SARS-CoV-1 nucleoprotein.

The secondary detection antibody was conjugated to a fluorophore which allows quantification of individual infected cells on a high throughput optical imaging system.

A minimum of 16,000 cells are counted per sample dilution across four wells - two each in duplicate plates. Data are reported based on a 4-parameter regression curve (using a constrained fit) as a 50% neutralization titer (IC50).

## Results

The ABO blood typing results from the initial 500 plasma units used for manufacturing hIVIG batches are presented in Table 1. Results from two published studies are included as comparators.

**Table 1:**
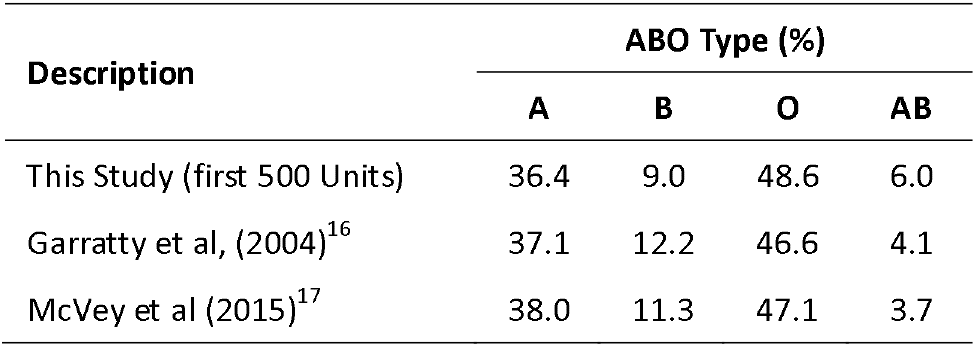
ABO Blood type distribution of convalescent plasma from test batches for this study compared to published values

| Description | ABO Type (%) |  |  |  |
| --- | --- | --- | --- | --- |
|  | A | B | O | AB |
| This Study (first 500 Units) | 36.4 | 9.0 | 48.6 | 6.0 |
| Garratty et al, (2004)<sup>16</sup> | 37.1 | 12.2 | 46.6 | 4.1 |
| McVey et al (2015)<sup>17</sup> | 38.0 | 11.3 | 47.1 | 3.7 |

To assess the recovery of the anti-SARS-CoV-2 specific antibodies, results from testing with an IgG specific ELISA and a neutralizing antibody assay are presented in Table 2 for the first product batches produced and their corresponding plasma pools. The IgG specific ELISA titers increased more than 10-fold when processing the pooled plasma into the final product. Neutralizing activity increased approximately three-fold. The IgG concentration was also increased by more than 10-fold from the pooled plasma to the final product. ELISA titers normalized to IgG were similar in the starting material and the finished product.

**Table 2:** Anti-SARS-CoV-2 titers and specific activities (n=3, ± standard deviation)

| Sample Point | ELISA Titer, 1:X | Neutralizing Activity Titer, IC50, 1:X | IgG, mg/mL | Normalized ELISA Titer, titer/mg /mL IgG |
| --- | --- | --- | --- | --- |
| Pool | 9038 ± 4037 | 110 ± 15 | 8.0 ± 0.2 | 1131 ± 497 |
| Final Product | 125089 ± 50905 | 325 ± 76 | 103 ± 3 | 1213 ± 477 |

Quality control test results from the first three batches are presented in Table 3. In these test batches, 100% of the protein content was IgG. The IgG was present entirely as monomers and dimers with aggregates and fragments below the limits of detection. A process impurity (sodium caprylate) and plasma protein impurities were found at very low concentrations in the final product.

**Table 3:** COVID-19 Hyperimmune Product Characteristics (n=3, ± standard deviation)

| Test | Average |
| --- | --- |
| **Potency** |  |
| ELISA Titer (1:X) | 125089 ± 50905 |
| ELISA Titer (ratio to control) | 27.1 ± 2.1 |
| Neutralization Titer (IC50, 1:X) | 325 ± 76 |
| **Formulation** |  |
| Glycine, M | 0.20 ± 0.02 |
| pH, diluted | 4.3 ± 0.1 |
| Protein Concentration, % | 10.3 ± 0.3 |
| Osmolality, mOsmol/kg | 246 ± 23 |
| **Molecular Profile and Purity** |  |
| Protein Composition, Gamma Globulin % | 100 |
| Monomer+Dimer, % | 100 |
| Aggregate, % | <1 |
| Fragment, % | <1 |
| Anti-Complement Activity, CH50/mg IgG | 0.47 ± 0.21 |
| Sodium Caprylate, µg/mL | <8 |
| IgA, mg/mL | 0.038 ± 0.003 |
| IgM, mg/mL | ≤0.007 |
| Prekallikrein Activator (PKA), IU/mL | <1.0 |
| FXIa, mIU/mL | <1 |
| Anti-A Titer, 1:X | 29.0 ± 5.2 |
| Anti-B Titer, 1:X | 16.0 |
| Anti-D Titer, 1:X | <2 |

## Discussion

ABO distribution in the plasma used to make these batches of hIVIG was found to be similar to published distributions.^16,17^ Modeling of the ABO distribution of units pooled was performed to control anti-A and anti-B levels, which are normally limited by dilution with larger batch sizes used in the manufacture of commercial immune globulin products. The modeled distribution resulted in products that all meet the anti-A and anti-B ≤1:64 requirements used by manufacturers of IGIV products to reduce the risk of hemolytic activity in patients.^18,19^ Hemolytic reactions are a rare event occurring in < 1 in 10,000 patients and are often without serious sequalae. These reactions are more common in non-O blood groups and have been found to be influenced by the blood type distribution of the plasma donors.^20^

Product characteristics for hIVIG were similar to those for Gamunex. The caprylate/chromatography purification process results in a highly purified IgG product that maintains the IgG activity found in the starting plasma. Additionally, experience with hyperimmune globulin preparations for rabies, tetanus and hepatitis B provide a reasonable expectation for the safety and effectiveness of hIVIG.

During processing, the normalized anti-SARS-Cov-2 antibody activity (expressed relative to IgG concentration) was maintained, as expected for a purified IgG product. Antibody neutralizing activity was increased approximately three-fold from the plasma pool to the final product. This increase in neutralizing activity indicates that patients treated with hIVIG compared to an equivalent volume of convalescent plasma would receive higher neutralizing activity. Alternatively, patients treated with hIVIG could receive a smaller treatment volume compared to treatment with convalescent plasma and potentially decrease the chances for transfusion-associated circulatory overload.

The three-fold increase in neutralizing activity from the pooled plasma to the final product, is smaller than expected increase based on ELISA titer. This may be caused by contributions from IgM and IgA which have been removed during purification of IgG.

IgM has been identified as a primary source of anti-A and anti-B intravascular hemolytic activity.^21^ IgM is removed to near or below the limits of detection from the hyperimmune product, hIVIG, greatly reducing the danger of this adverse event. In contrast, when patients are treated with convalescent plasma, they must be matched by donor blood type to reduce the chances of hemolysis.

Similarly, removal of IgA provides a potential therapeutic advantage for hIVIG over convalescent plasma in patients who are IgA deficient and may have been previously treated with blood products and formed antibodies to IgA. Although purified IgG products contain trace amounts of IgA and are labeled to warn for potential adverse events, reactions in IgA deficient patients are extremely rare ^22^.

A potential advantage hIVIG over direct administration of plasma from individuals or a monoclonal antibody is the diversity of antibodies obtained from a pool of convalescent donors which may provide a wider range of anti-viral activity.^23^ This diversity may be important in overcoming viral mutations, although thus far, the viral genome has remained relatively stable.^24^ Antibody diversity may provide a broader range of anti-viral activity by attacking different viral epitopes and enlisting different cellular mechanisms.

In summary, the processes described here describe the manufacturing of multiple batches of anti-SARS-CoV-2 hyperimmune globulin (hIVIG) that met regulatory requirements. Clinical trials to study the efficacy of hIVIG have been initiated. The first clinical study in which the hIVIG is being evaluated is the Inpatient Treatment with Anti-Coronavirus Immunoglobulin (ITAC) trial.^25^

## Acknowledgements

The authors thank the NIAID Integrated Research Facility, Frederick, Maryland, for antibody neutralization testing. Michael K. James, Ph.D. and Jordi Bozzo, Ph.D. are acknowledged for writing and editorial assistance.

## References

1. Huang C, Wang Y, Li X, et al. Clinical features of patients infected with 2019 novel coronavirus in Wuhan, China. The Lancet 2020;395: 497–506.

2. Timeline of WHO’s response to COVID-19: Last updated 9 September 2020 [monograph on the internet]. Geneva: World Health Organization; 2020. Available from: https://www.who.int/news-room/detail/29-06-2020-covidtimeline

3. Shen C, Wang Z, Zhao F, et al. Treatment of 5 Critically Ill Patients With COVID-19 With Convalescent Plasma. JAMA 2020;323: 1582–9.

4. Luke TC, Kilbane EM, Jackson JL, et al. Meta-Analysis: Convalescent Blood Products for Spanish Influenza Pneumonia: A Future H5N1 Treatment? Ann Intern Med 2006;145: 599–609.

5. Soo YO, Cheng Y, Wong R, et al. Retrospective comparison of convalescent plasma with continuing high-dose methylprednisolone treatment in SARS patients. Clin Microbiol Infect 2004;10: 676–8.

6. Ko JH, Seok H, Cho SY, et al. Challenges of convalescent plasma infusion therapy in Middle East respiratory coronavirus infection: a single centre experience. Antivir Ther 2018;23: 617–22.

7. Jungbauer C, Weseslindtner L, Weidner L, et al. Characterization of 100 sequential SARS-CoV-2 convalescent plasma donations. Transfusion 2020.

8. 21 CFR 630 - General requirements for blood and blood components intended for transfusion or further manufacturing use. In: Code of Federal Regualtions, US Department of Health and Human Services, Washington, D.C.: United States Government Printing Office, 2010.

9. Plasma Protein Therapeutics Association: Safety and quality [monograph on the internet]. 2020. Available from: https://www.pptaglobal.org/safetv-qualitv

10. Investigational COVID-19 Convalescent Plasma: Guidance for Industry. Bethesda, MD: United States Food and Drug Administration, Department of Health and Human Services; Updated September 2, 2020. Available from: https://www.fda.gov/media/136798/download

11. Lebing W, Remington KM, Schreiner C, et al. Properties of a new intravenous immunoglobulin (IGIV-C, 10%) produced by virus inactivation with caprylate and column chromatography. Vox Sang 2003;84: 193–201.

12. Alonso W, Vandeberg P, Lang J, et al. Immune globulin subcutaneous, human 20% solution (Xembify(R)), a new high concentration immunoglobulin product for subcutaneous administration. Biologicals 2020;64: 34–40.

13. Gamunex-C [Immune Globulin Injection (Human) 10% Caprylate/Chromatography Purified]-Package Insert [monograph on the internet]. Research Triangle Park, NC: Grifols Therapeutics LLC; 2020. Available from: https://www.gamunex-c.com/documents/27482625/27482925/Gamunex-C+Prescribing+Information.pdf/9258bd0f-4205-47e1-ab80-540304c1ff8e

14. Chapter 151: Pyrogen test. In: United States Pharmacopeia 43 – National Formulary 38, p. 6577 (2020).

15. Chapter 71: Sterility tests. In: United States Pharmacopeia 43 – National Formulary 38, p. 6481 (2020).

16. Garratty G, Glynn SA, McEntire R. ABO and Rh(D) phenotype frequencies of different racial/ethnic groups in the United States. Transfusion 2004;44: 703–6.

17. McVey J, Baker D, Parti R, et al. Anti-A and anti-B titers in donor plasma, plasma pools, and immunoglobulin final products. Transfusion 2015;55 Suppl 2: S98–104.

18. Monograph 0918: Human normal immunoglobulin for intravenous administration. In: European Pharmacopeia 10.0, Strasbourg, France: Council of Europe, pp. 2862–2863 (2019).

19. General Chapter 2.6.20: Anti-A and anti-B haemagglutinins (indirect method). In: European Pharmacopeia 10.0, Strasbourg, France: Council of Europe, pp. 218–219 (2019).

20. Sewell WAC, Kerr J, Behr-Gross ME, et al. European consensus proposal for immunoglobulin therapies. Eur J Immunol 2014;44: 2207–14.

21. Flegel WA. Pathogenesis and mechanisms of antibody-mediated hemolysis. Transfusion 2015;55 Suppl 2: S47–58.

22. Pineda AA, Taswell HF. Transfusion Reactions Associated with Anti-IgA Antibodies: Report of Four Cases and Review of the Literature. Transfusion 1975;15: 10–5.

23. Liu L, Wang P, Nair MS, et al. Potent neutralizing antibodies against multiple epitopes on SARS-CoV-2 spike. Nature 2020;584: 450–6.

24. van Dorp L, Acman M, Richard D, et al. Emergence of genomic diversity and recurrent mutations in SARS-CoV-2. Infect Genet Evol 2020;83: 104351.

25. Inpatient treatment with anti-coronavirus immunoglobulin (ITAC) [monograph on the internet]. clinicaltrials.gov; 2020. Available from: https://clinicaltrials.gov/ct2/show/NCT04546581

